# Epistasis meets pleiotropy in shaping biophysical protein subspaces associated with antimicrobial resistance

**DOI:** 10.1101/2023.04.09.535490

**Authors:** C. Brandon Ogbunugafor, Rafael F. Guerrero, Eugene I. Shakhnovich, Matthew D. Shoulders

**Affiliations:** Department of Ecology and Evolutionary Biology, Yale University, New Haven, CT; Department of Chemistry, Massachusetts Institute of Technology, Cambridge, MA; Santa Fe Institute, Santa Fe, NM; Department of Biological Sciences, North Carolina State University, Raleigh, NC; Department of Chemistry and Chemical Biology, Harvard University, Cambridge MA

**Author notes:** **Corresponding author:** C. Brandon Ogbunugafor.

**Keywords:** epistasis, pleiotropy, protein space, fitness landscape, biophysics, antimicrobial resistance

## Abstract

Protein space is a rich analogy for genotype-phenotype maps, where amino acid sequence is organized into a high-dimensional space that highlights the connectivity between protein variants. It is a useful abstraction for understanding the process of evolution, and for efforts to engineer proteins towards desirable phenotypes. Few framings of protein space consider how higher-level protein phenotypes can be described in terms of their biophysical dimensions, nor do they rigorously interrogate how forces like epistasis—describing the nonlinear interaction between mutations and their phenotypic consequences—manifest across these dimensions. In this study, we deconstruct a low-dimensional protein space of a bacterial enzyme (dihydrofolate reductase; DHFR) into “subspaces” corresponding to a set of kinetic and thermodynamic traits [(*k_cat_, K_M_*, *K_i_*, and *T_m_* (melting temperature)]. We then examine how three mutations (eight alleles in total) display pleiotropy in their interactions across these subspaces. We extend this approach to examine protein spaces across three orthologous DHFR enzymes (*Escherichia coli, Listeria grayi*, and *Chlamydia muridarum*), adding a genotypic context dimension through which epistasis occurs across subspaces. In doing so, we reveal that protein space is a deceptively complex notion, and that the process of protein evolution and engineering should consider how interactions between amino acid substitutions manifest across different phenotypic subspaces.

## 1. Introduction

For all the sophistication of technologies associated with studying protein structure function—cryoEM, AlphaFold and deep mutational scanning, for example—basic questions remain about how we consider and measure the shape of genotype-phenotype maps in the study of proteins. Addressing these questions requires theoretical and conceptual instruments that can be used to understand how genotype and amino acid composition confers phenotype at the protein level. In this regard, evolutionary biologists have used two somewhat related analogies—the fitness landscape and protein space—to describe how evolution searches through the space of possibility from genotype to protein phenotype [1–7].

One important conceptual innovation in the study of genotype-phenotype maps involves the notion that phenotypes can often be deconstructed into “micro-landscapes” that are parts of a larger fitness landscape [8]. In the case of some enzymes, there are multiple genotype-phenotype maps corresponding to different biophysical phenotypes: for example, those related to enzyme kinetics and those defining thermodynamic properties. A foundational study in this area (2005) reconstructed a protein fitness landscape associated with the use of a co-enzyme from component biophysical traits [9]. In more recent examples, traits like the IC_50_ or minimal inhibitory concentration (both metrics for drug resistance) when applied to enzyme targets of drugs (like dihydrofolate reductase or beta-lactamase), can be similarly deconstructed into landscapes corresponding to kinetic and thermodynamic parameters [8, 10]. In the language of protein space, the hierarchy between complex phenotypes and component phenotypes can be framed in terms of “subspaces.”

If we consider the possibility that protein space is composed of biophysical subspaces, new questions arise surrounding how those subspaces are constructed, and what their existence and shape means for protein evolution. For example, epistasis—defined colloquially as the “surprise at the phenotype when mutations are combined, given the constituent mutations’ individual effects” [11] has long been known to craft the topography of fitness landscapes. Epistasis remains a provocative concept because it shapes genotype-phenotype maps in surprising ways, creating rugged fitness landscapes with fitness valleys that can undermine or constrain the process of adaptation [12–21]. If the protein space through which evolution is operating is composed of several subspaces (Figure 1), then one might ask how epistasis manifests across these subspaces.

**Figure 1:**
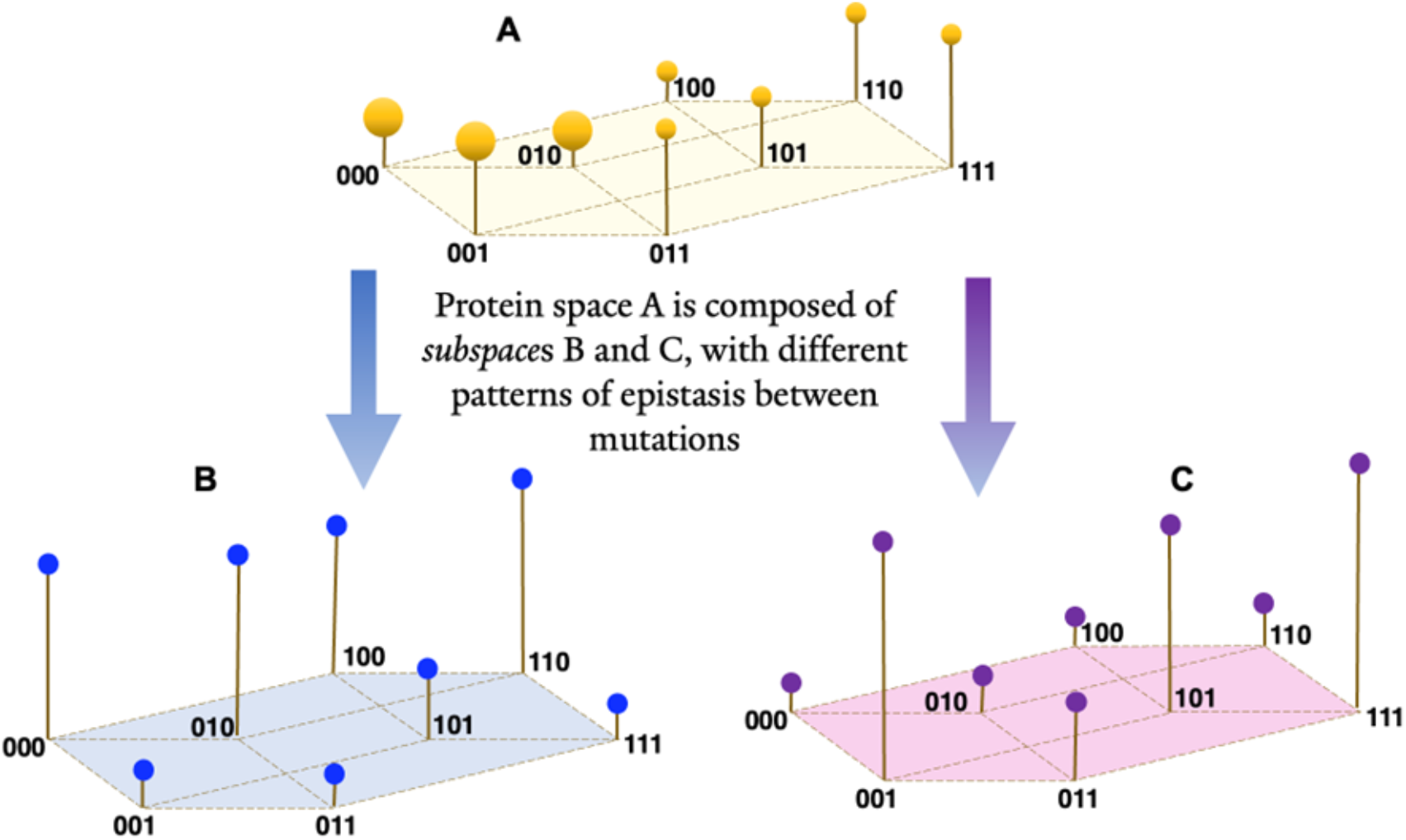
A 3-D conceptual diagram of protein space and subspace, revealing how different epistatic interactions manifest across different traits (pleiotropy). This image outlines the central concept behind the complex structure of protein-level phenotypes, organized into protein space. The complex phenotype (A) can be deconstructed into subspaces (B) and (C). The heights of the lines correspond to a phenotype value for that particular space-trait. Note the differences in topography. Importantly, these differences manifest because of differences in epistasis between mutations. The [0] and [1] values correspond to the presence (1) or absence (0) of a mutation at a given location. The size of the protein spaces in this schematic (including eight alleles) is the same as that used in this study, but this need not be the case. The subspace concept transcends a space of any size—one could construct the space-subspace dichotomy for spaces of hundreds or thousands of nodes.

In examining how mutation effects manifest across different subspaces, we run into a different (perhaps equally) provocative concept from evolutionary theory: pleiotropy, which can be defined by the differential effects of genes or mutations on seemingly disparate traits [22]. Pleiotropy is frequently used in discussions around tradeoffs in evolution, as in a presumed tradeoff between generalism and specialism [23] or pathogen virulence and transmission [24]. However, the concept has a greater reach: it forces us to reconsider the phenotypic effects of every allele or mutation, as the effects that we focus on might not be the only (or most meaningful) trait affected.

Relatedly, several studies have examined how nonlinear interactions between loci in a gene (and their putative amino acids) manifest across the traits that correspond to subspaces [8–10, 25–27]. This topic continues to be an important area of examination, because while protein evolution may be tracked using high-level phenotypes (e.g., fitness, IC_50_ or minimal inhibitory concentration), evolution is often operating incongruously across the different subspaces (e.g., thermodynamic versus kinetic components). Consequently, focusing on the shape of these subspaces—dictated by epistasis—is important for resolving and predicting the phenotypic effects of mutations and their putative amino acid substitutions (as in protein evolution and bioengineering).

In this study, we deconstruct sets of protein spaces (composed of eight alleles) for DHFR orthologs in *Escherichia coli, Listeria grayi*, and *Chlamydia muridaum* into subspaces corresponding to different biophysical traits: *k_cat_, K_M_*, *K_i_*, and *T_m_* (melting temperature). In doing so, we reveal how the shapes of protein subspaces can differ profoundly according to biophysical trait, and quantify the epistasis operating in each of the subspaces (corresponding to biophysical trait). Finally, we discuss the implications of this in the present and future study of evolutionary theory, protein evolution and bioengineering.

## 2. Methods

### 2.1. Laboratory measurement of biophysical traits

The data in this study originated from a prior study of the biophysical decomposition of a fitness landscape [8]. We have, however, provided the data generated from that study in the Supplemental Information. The methods for measuring the biophysical traits in this study were also previously described in a prior study [8, 28]. We refer authors interested in replicating the laboratory-derived biochemical and biophysical to those studies.

### 2.2. Nomenclature

For translation purposes, we will employ a particular nomenclature for discussing the mutations and DHFR alleles used in this study. The mutations corresponding to P21L, A26T, and L28R in *E. coli* and *L. grayi* are referred to with regards to their combinatorial arrangement. For example, “PAL” corresponds to the enzyme variant with amino acids Proline (P), Alanine (A), and Leucine (L) at the three loci of interest. In *C. muridarum*, the orthologous mutations are P23L, E28T and L30R.

### 2.3. Notes on the bacterial orthologs

In Supplemental Table 1, we observe the sequence identity matrix for DHFRs derived from *E. coli, L. grayi*, and *C. muridarum*. These data are also present in a prior study that examined this same suite of enzymes [8].

### 2.4. Comparing the topography of the protein spaces/fitness landscapes

. The protein spaces were constructed from existing data (see [29] and Supplemental Information), and were then compared using the Kendall rank order test and matrix. This test measures the concordance or discordance of the landscapes with respect to the order of the alleles in a landscape. For each species, we ranked genotypes within subspace assigning values from 1 (maximum) to 8 (minimum value for the measurement in that subspace). We used these ranks to calculate rank correlations (Kendall’s *τ*) across all subspace pairs (a method implemented in R; base and corrr packages; [30]).

### 2.5. Higher-order epistasis

The system explored in this study has been previously examined with respect to how proteostasis machinery influences higher-order epistasis [27, 31]. And while other studies have measured epistasis on biophysical traits [8, 10], few have rigorously examined and compared how higher-order epistasis manifest across subspaces, or directly compared the shapes of these spaces across species orthologs.

To measure epistasis, we use a method developed in theoretical computer science termed the Walsh-Hadamard transform, which computes a coefficient corresponding to the magnitude and sign of interaction between mutations, akin to an epistatic coefficient. It was pioneered for use in the study of epistasis in a 2013 study that both provided a primer for the calculation and analyzed several combinatorically complete data sets [11]. The Walsh-Hadamard transform has since been further elaborated on and applied to the study of higher-order epistasis across an array of empirical data sets [32–34].

While we will describe certain features of the analysis here, we suggest that those interested in further details refer to existing manuscripts that describe and apply the method to datasets similar in structure to the ones analyzed here [11, 32]. Note that this approach is only one of myriad methods that one can use to quantify epistasis, and we encourage those interested to engage several reviews that have addressed this topic directly [21, 26, 35, 36]. Moreover, there are new methods that facilitate the measurement of epistasis in large genomic data sets [37, 38].

The Walsh-Hadamard transform implements phenotypic measurements into a vector, then a Hadamard matrix, subsequently scaled by a diagonal matrix. The calculation yields a set of coefficients that measure the degree to which the relationship between genetic information and phenotypes are linear, or second order, third, and so forth.

One limitation of the Walsh-Hadamard transform is that the data it employs are generally combinatorially complete with no more than two variants at a given locus of information. More recent studies have, however, proposed strategies to transcend some of the presumed limitations [39]. Nonetheless, in this study we utilize the methods on a combinatorially complete set of mutants, where we can represent the presence or absence of a given mutation by a 0 or 1 at a given site. For example, we can represent a wild-type variant of a gene as 000. In this scenario, mutations at each of three sites (e.g., the three mutations corresponding to trimethoprim resistance in *E. coli* dihydrofolate reductase P21L, A26T, and L28R) encoded as 111 (For example, see Figure 1).

The full data set for the alleles consists of a vector of phenotypic values (resistance to trimethoprim in the case of the DHFR mutants) for all possible combinations of mutations (8 in total), represented by their single amino acid substitutions:

For *E. coli* and *L. grayi*:

PAL, LAL, PAR, PTL, LAL, PTR, LAR, LTL, LTR

For *C. muridarum*:

PEL, LEL, PER, PTL, LEL, PTR, LER, LTL, LTR

In binary notation, where [0] is the wild-type genotype and the [1] the mutant, we can convert the above into a different notation:

000, 100, 001, 010, 100, 011, 101, 110, 111

This vector of genotypes can be multiplied by a (8 x 8) square matrix, which is the product of a diagonal matrix *V* and a Hadamard matrix *H*. These are defined recursively by:

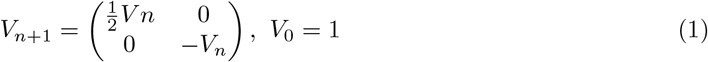

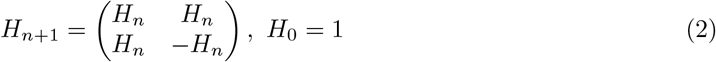

where n is the number of mutations that define protein spaces in the DHFR orthologs (*n* = 3 in this study). The multiplication gives the following expression:

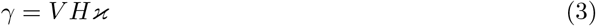

H and V are the matrices from equations 1 and 2 and *γ* is the Walsh coefficient, the measure of the average epistatic interaction between amino acid substitutions in the DHFR orthologs:

Positive values would correspond to a positive effect on a given biophysical phenotype. Negative values for an effect suggest that the average epistatic effect is negative on the biophysical trait.

### 2.6. Determining higher-order interactions across biophysical trait

The above formula can be used to calculate the strength of interactions between parcels of information, that is, the amino acids corresponding to the mutations in DHFR in our example. But what about higher-order interactions (epistasis beyond pairwise in this case)?

Previous studies have examined how higher-order epistasis manifests in adaptive landscapes that include analogously structured data sets, including other enzymes [8, 11, 20, 34]. In a complete data set comprising eight variants, we can describe the *interactions* between individual loci and genetic background in binary terms. If we are talking about a combinatorial set of variants with three loci, we can describe the *interactions* using binary representation.

This is a key point of clarity: unlike many studies of fitness landscapes that use binary notation to describe the presence or one mutation or the other, here we use it strictly to describe the phenotypic effect of a given mutation *averaged across all possible single mutation background combinations*:

000: The effect of the interaction between the mutations in the wild-type background.
**1: The average phenotypic effect of the interaction between the “third site” mutation (L28R in *E. coli* and *L. grayi*; L30R in *C. muridarum*) and the other set of possible individual mutations in the set of eight.
*1*: The average phenotypic effect of the interaction between the “second site” mutation, A26T in *E. coli* and *L. grayi*; E28T in *C. muridarum*) and all the other set of possible individual mutations in the set of eight.
1**: The average phenotypic effect “first site” (P21L in *E. coli* and *L. grayi*; P23L in *C. muridarum*) mutation and all possible individual mutations in the set of eight.
*11: The average phenotypic effect of the pairwise (second-order) interaction between mutations at the second and third loci and all the other set of possible individual mutations in the set of eight.
1*1: The average phenotypic effect of the pairwise (second-order) interaction between mutations at the second and third loci and all the other set of possible individual mutations in the set of eight.
11*: The average phenotypic effect of the pairwise (second-order) interaction between mutations at the first and second loci and all the other set of possible individual mutations in the set of eight.
111: The average phenotypic effect of the third-order interaction between mutations at all three loci and all the other set of possible individual mutations in the set of eight.

In the set of interactions that we measure, there is one zeroth order effect, three first-order interactions, three second-order interactions, and one third-order interaction. The third-order interaction would formally qualify as “higher-order.”

In addition, one can take the mean of these epistatic coefficients within an order, which can facilitate comparisons between orders. For a given epistatic coefficient we compute an epistatic coefficient, E, as in prior studies that have examined higher-order interactions on empirical fitness landscapes [34].

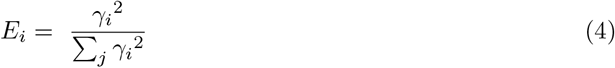

Though we have used absolute values in prior examinations of epistasis (e.g., [34]), this iteration of the calculation utilizes the squares, as it is more synonymous with physics interpretations of signal strength (an appropriate analogy for understanding the strength of interactions in epistasis). This calculation translates to the intensity of the interactions corresponding to a specific order. For example, 1^*st*^ order effects to main effects, *2^nd^* order to pairwise effects, and *3^rd^* order to three-way effects.

### 2.7. A brief note on the language of protein space vs. adaptive or fitness landscape

. Some of the concepts explored here have previously been framed in terms of protein fitness landscapes [6, 40]. The concepts of protein space and the fitness landscape are at least compatible, even identical in some cases. And previous work has explored the similarity between protein space and the fitness landscape [41–43]. For many problems in molecular evolution, both can be used. But the subtle differences in their framing are important to articulate here.:

1. The fitness or adaptive landscape analogy is most appropriate as a representation of genotypephenotype maps when examining an evolving population. Protein space, on the other hand, is less fixated on any solution or “fitness peak,” but rather focuses on the broader notion that evolutionary possibility can be mapped across an *n*-dimensional space.
2. Relatedly, the fitness or adaptive landscape concept can be encumbered by the definition of “fitness.” Protein space can describe the relationship between mutational neighbors (nodes in the space) with respect to any conceivable phenotype, whether it be adaptive or not.

## 3. Results

This study aimed to examine the topography of biophysical protein subspaces across three orthologs of DHFR (*E. coli, L. grayi*, and *C. muridarum*). We offer that pairwise and higher-order epistasis are the driving engine behind the observed topographical differences across orthoglos and subspaces. Our findings are organized and described in three major categories:

- Comparisons between the topography of protein subspaces of DHFR across biophysical traits and bacterial species.
- Measuring the epistasis between loci.
- Comparison of the higher-order epistasis that drives these differences in subspace topography.

Where relevant, we will discuss statistical tests used, our rationale, and the conclusions from those analyses.

### Comparing the topography of the subspaces

We first organized the data into independent subspaces. We depict the resulting subspaces in terms of how the scaled values (shown in standard deviations) change across biophysical traits (Figure 2). We then used a Kendall rank order correlation to compare the topographies of the various landscapes (see Methods). In Figure 3, we observe that many of the significant findings involve the kinetic traits *K_i_, k_cat_* and *K_M_*. For example, focusing just on the *E. coli* subspaces: there is a strong negative correlation between *K_i_* and *k_c_at* (*p* < 0.01). Also for *E. coli*, note the less strong but significant concordance between *k_cat_* and *K_M_* (*p* < 0.05). Similar patterns can be observed within and between the other species, reflecting both biophysical rhyme and reason (again, the kinetic traits are related), and widespread variation across the expanse of protein spaces.

**Figure 2:**
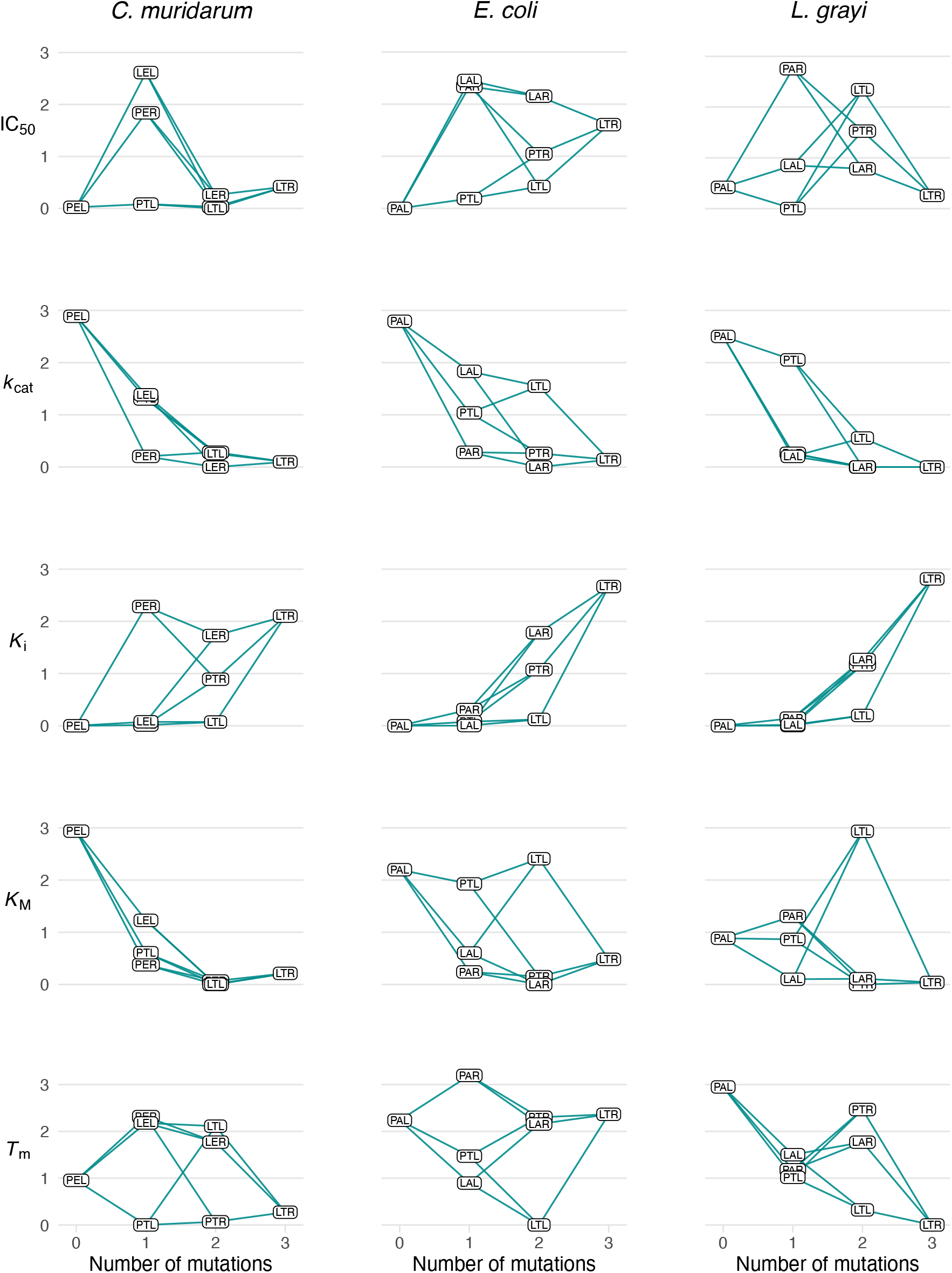
Measurements of protein subspaces across orthologous bacterial enzymes (DHFR). Scaled values are shown in standard deviations. Measurements of the different subspaces for the suite of mutations associated with resistance to antifolate drugs in dihydrofolate reductase. Values for each subspace are scaled according to the mean of the value for that trait. Comparing the topography of the subspaces within and across species suggests varying patterns of similarities and differences across subspaces.

**Figure 3:**
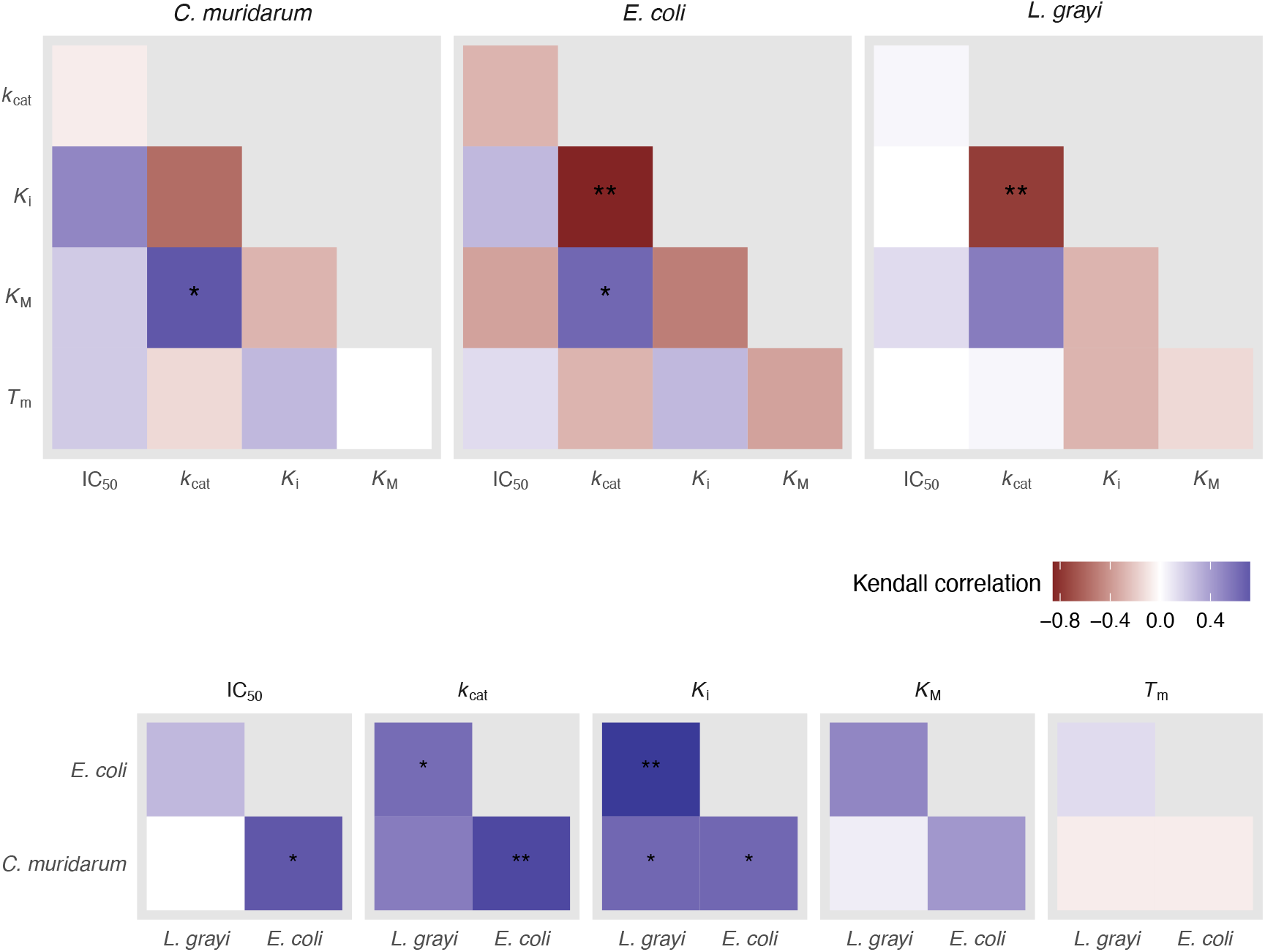
A comparison of topography across subspaces. We use a Kendall rank-order test to quantify correlations among the landscapes of five traits in three species. * = p < 0.05, ** = p < 0.01. Concordant and discordant subspaces tend to be focused on the kinetic traits: *K_M_*, *k_c_at*, and *K_i_*. See Figure S1 for a depiction of the rank orders.

Next, we computed these epistatic effects as outlined in the Methods (Figure 4), even ranking the interactions and effects, across trait and bacterial orthologs. We then re-organized the effects from Figure 5 into higher-order terms. That is, we squared the effects depicted in Figure **??** and summed them according to order—zeroth, first, second (pairwise), or third (see Methods). This lens highlights that the manner in which mutations are interacting differs drastically across subspace trait, and across orthologs.

**Figure 4:**
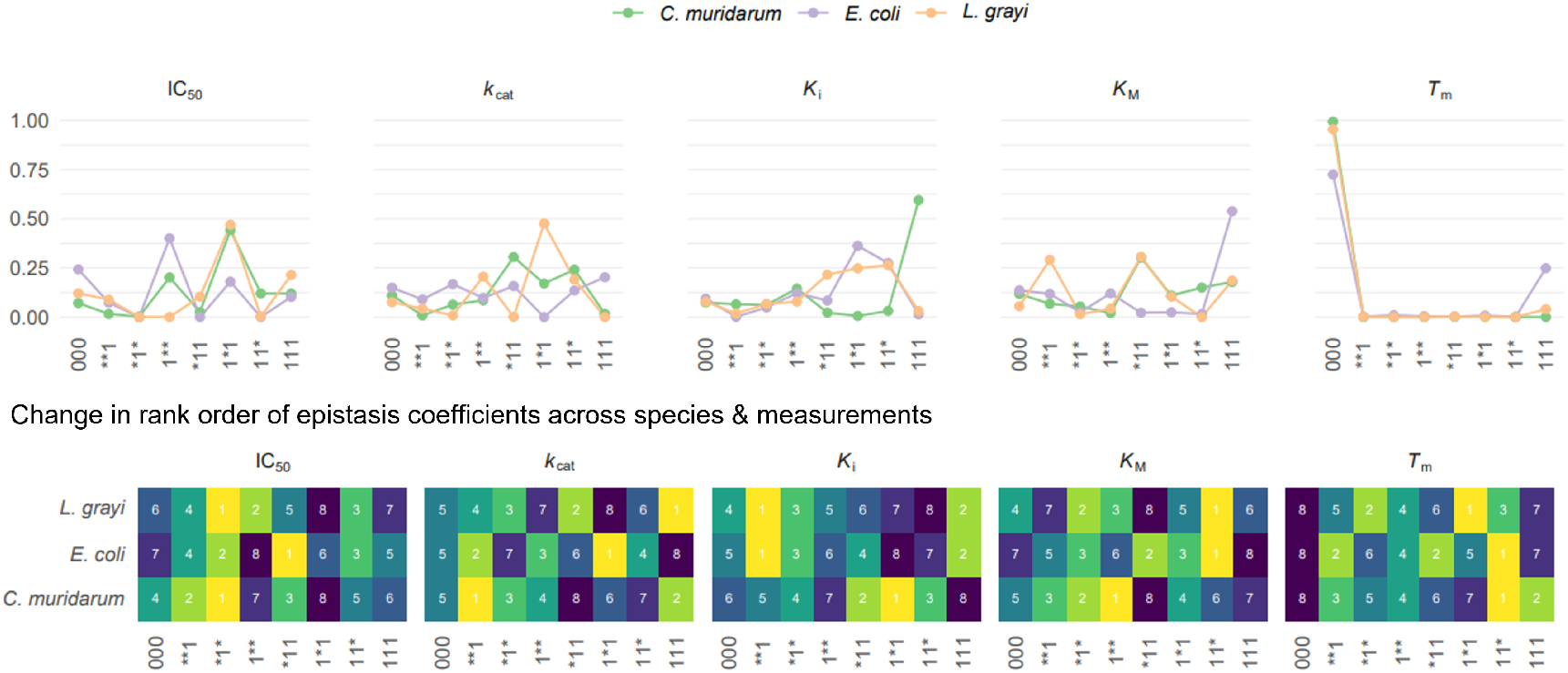
Epistasis meets pleiotropy: measurements of epistasis across orthologous subspaces. We compute the Walsh-Hadamard coefficient for each sort of interaction. Top panel: Individual graphs correspond to different traits, starting with the complex trait (IC_50_) on the left, followed by the individual biochemical and biophysical traits (*K_i_*, *K_M_*, *k_cat_*, and *T_m_*). X-axis depicts individual mutation effects, with [1]s corresponding to the presence of a mutation at a given location. A [1] at the first site corresponds to the presence of the P21L (*E. coli* and *L. grayi*) or P23L mutation (*C. muridarum*). A [1] at the second site corresponds to the A26T (*E. coli* and *L. grayi*) or E28T (*C. muridarum*) mutations, and a [1] at the last site corresponds to the L28R (*E. coli* and *L. grayi*) or L30R (*C. muridarum*) mutations. Bottom panel: The same data as in the top panel, depicted as rank orders of effects. For example, for *L. grayi* the 1*1 pairwise interaction between the P21L and A26T mutations is the highest ranked (has the highest magnitude) of all of the interactions.

**Figure 5:**
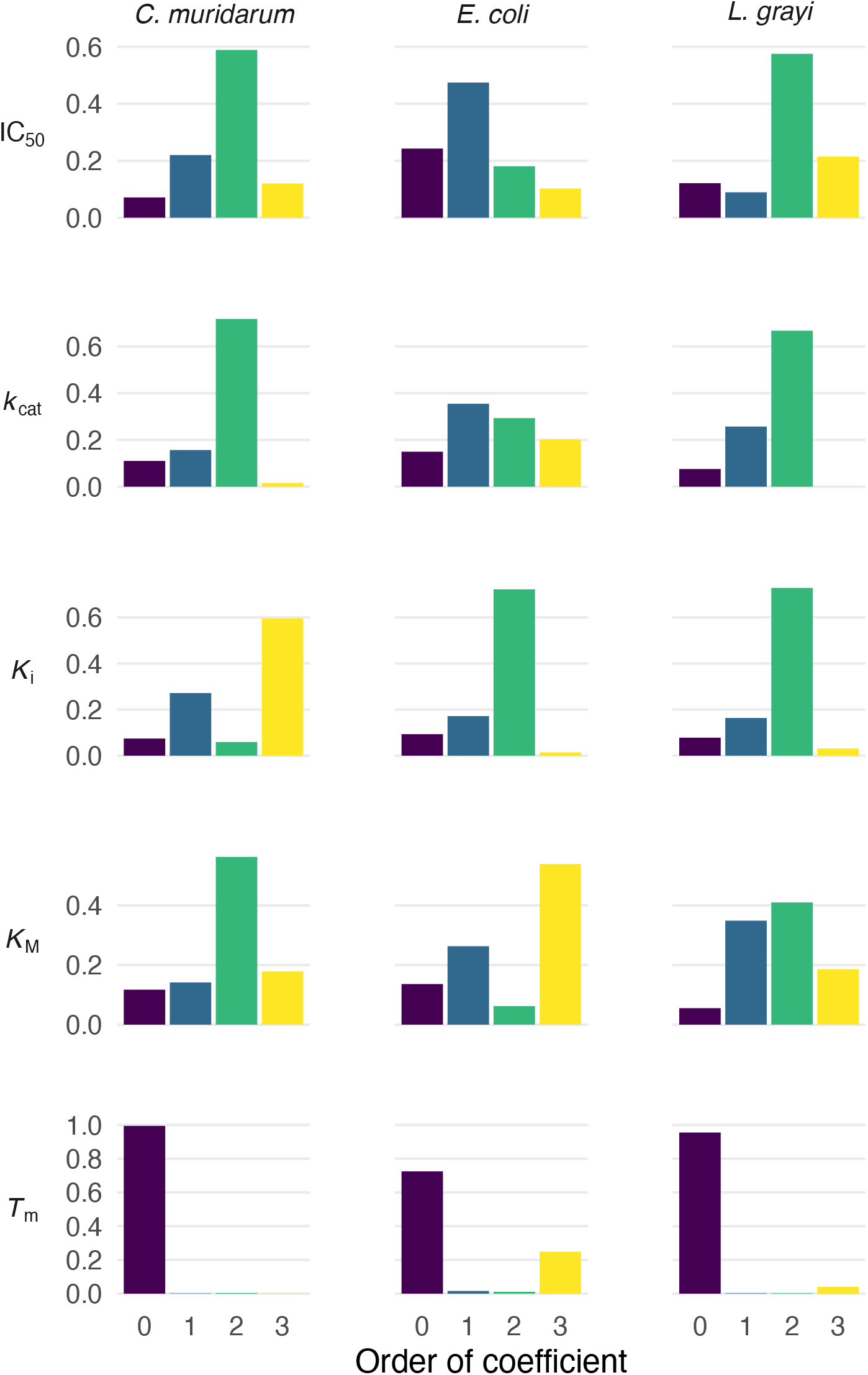
Epistasis meets pleiotropy, organized by order, across orthologous subspaces: As outlined in the methods, the epistatic coefficients can be reorganized to depict the effects by order. This approach aids in efforts to compare the overall presence of epistatic effects of a certain order across biophysical subspaces. We observe that different sorts of interactions govern certain biophysical traits. For example, *K_M_* is dominated by pairwise interactions in *C. muridarum*, third-order interactions in *E. coli*, and first-order and pairwise interactions in *L. grayi*.

For *E. coli*, pairwise and third-order effects predominate in *K_M_* and *K_i_. T_m_*, however, is most influenced by the combination of wild-type mutations. For *L. grayi*, pairwise effects dominate the kinetic traits — *K_M_*, *k_cat_*, and *K_i_* — with third order effects playing a meaningful role in abundance. Recall from the comparison of the topography of landscapes (Figure 3) that these traits were the ones with the strongest patterns of concordance or discordance. While not a rule that applies across the entire data set, this theme suggests that similar patterns of epistasis operate on traits that are related to one another biologically, a finding that is consistent with our intuition.

Zeroth order effects (corresponding to the combination of mutations present in the wild type *L. grayi* DHFR) are especially meaningful in the *T_m_* subspace. In *C. muridarum*, pairwise or third order effects dominate every subspace except for *T_m_* subspace, where zeroth order effects predominate. Indeed, while patterns differ wildly across ortholog subspaces, one consistent observation is the relative lack of higher-order effects operating on the *T_m_* subspace. In all three species, zeroth order effects were a notable influence on *T_m_*.

### Analysis of epistasis that underlies differences in subspace topography

Using the Walsh-Hadamard Transform, we then calculated the average effects of mutations, as well as the pairwise and three-way effects, on a range of traits across the IC_50_ and biophysical subspaces. These calculations revealed large differences in patterns of epistasis across subspace traits (Figure 4). Note again that IC_50_ is depicted alongside the subspaces, for visualization purposes, so that we can see how the subspaces compare to the overarching space.

## 4. Discussion

In this study, we measured epistasis across four biophysical subspaces (*k_cat_, K_M_*, *K_i_*, and *T_m_*) of three orthologs (*Escherichia coli*, *Listeria grayi*, and *Chlamydia muridarum*) of dihydrofolate reductase, an enzyme target of antimicrobial drugs. Our findings fortify the notion that epistatic interactions remain a major challenge in resolving phenotype from genotype, because mutation effect interactions can tune the shape subspaces of a single protein differently. In this way, our study offers insight into the interface between two population genetics concepts—epistasis and pleiotropy—each of which are important forces in the process of adaptive evolution [44, 45].

We observe that the shape of protein space differs across orthologs of DHFR (Figures 2 – 4). This finding is unsurprising and emphasizes how even relatively minor differences in amino acid sequence (corresponding to the three species of bacteria) can have meaningful consequences for how protein space is structured. Figures 2 and 3 highlight several relationships (both concordance and discordance) between the shapes of protein spaces associated with IC_50_, *k_cat_, K_M_*, and *K_i_*. For example, there is strong discordance between *k_cat_* and *K_i_* across the protein spaces, and relatively strong concordance between *k_cat_* and *K_M_*. These are intuitive, as they are subspace phenotypes that are properties of the enzyme active site, where both Michaelis-Menten and mechanistic expectations for these results (e.g., that *k_cat_*, and *K_i_* should be discordant). By contrast, note that the *T_m_* protein subspaces (composed of 16 alleles) appear to be uncorrelated–neither concordant nor discordant–across the three species of bacteria.

Measures of epistasis (Figure 4 and 5) tell an important part of the story. While there are patterns of epistatic interactions across orthologs and traits, there are no universal properties (e.g., that a given order of interactions dominates a subspace trait). Alternatively, each subspace has patterns of similarity and difference according to trait and ortholog. For example, pairwise interactions between mutations appear to be an actor in many kinetic traits, but third-order interactions are relatively low in magnitude in the *L. grayi* ortholog of DHFR, across traits. Notably, the *T_m_* subspace is dominated by zeroth order effects—where mutations in the wild-type genotype had the largest interaction effect (Figure 5). This observation complements the comparisons of concordance or discordance depicted in Figure 3 (and discussed above), and could be related to the thermodynamics that contribute to a given protein’s *T_m_*, which may be more reliant on global features of an amino acid sequence, rather than peculiar interactions between mutations that influence resistance to a small molecular. Future studies will examine this at a more rigorous level.

### 4.1. Study limitations

This study has several limitations. The protein spaces explored are low-dimensional, each composed of only eight nodes. This represents a very small slice of the true protein space (astronomical in size), encompassing only a set of engineered mutations corresponding to those identified in experimental populations of bacteria exposed to trimethoprim [46]. Furthermore, the conversation about epistasis in evolutionary theory has grown in sophistication in recent years, with ideas such as “global epistasis” adding a new point of intrigue. Global epistasis refers to the notion that epistatic effects follow a system-wide pattern (e.g., diminishing returns) and arise from linear relationships between the phenotypic effects of a mutation and the fitness of the genetic background [47–50]. Global epistasis, as a phenomenon of non-linear genotype-to-phenotype mapping, is likely to compound the effects we describe in our study. The genotype-to-phenotype map is specific to each trait, so global “diminishing returns” effects are likely to affect each subspace differently. This constitutes a current area of investigation.

### 4.2. Ideas, speculation, and future directions

Our results have direct implications for efforts to engineer proteins using directed evolution or other approaches. For example, evolving a thermostable enzyme would amount to selection across one of the subspaces measured in our study (*T_m_*). Our study suggests that such directed evolution efforts should not only consider how mutations associated with increased thermostability interact epistatically but also the pleiotropic consequences of this epistasis^1^ on other traits. Importantly, our study differs from a foundational 2005 study that examined epistasis on component biophysical traits of an enzyme (isopropylmalate dehydrogenase; IMDH) [9]. In that study, epistasis was minimal across biophysical traits, but manifested at the higher level of protein fitness. Our study shows that not only is epistasis acting on biophysical traits, but these patterns also differ from subspace trait to subspace trait. There is no obvious explanation for why there was no apparent epistasis in the biophysical subspaces for IMDH evolving differential co-enzyme use, but there was substantial epistasis in our study of DHFR with evolved differences in resistance to an antifolate drug. That different enzymes have unique patterns of epistasis is unsurprising: they are different in structure and function. Our findings emphasize the importance of not over-generalizing results from study of a single, or just a few, enzymes.

Further, future studies should utilize newer technologies to examine protein space at a larger scale. For example, the use of deep mutational scanning has revealed substitutions in SARS-CoV-2 proteins that may be relevant for the design of vaccines and therapeutics [51–53], and revealed how host cell chaperones shape the evolution of viral pathogens [54–56]. These tools may reveal how epistasis and pleiotropy play out across subspaces that are thousands of nodes in size.

### 4.3. Conclusion

Future efforts to direct the evolution of protein phenotypes might be improved with the knowledge of how subspaces function as biophysical and biochemical “knobs” that tune higher-level protein phenotypes. Further, in epidemic settings, the effects of mutations on larger-scale pathogen phenotypes such as transmissibility (e.g., SARS-CoV-2) might be better diagnosed and understood mechanistically through examination of their subspace effects.

Our findings suggest that even single-locus complex traits —like the IC_50_ of an enzyme target of drugs—contain biophysical multitudes. These subspace multitudes have several implications for how we consider the problem of proteins searching space during evolution. Rather than evolution “up” or “across” a rugged fitness landscape, protein evolution or engineering can be more readily described as some combination of searches through different subspaces.

## Acknowledgments

The authors thank M. Miller-Dickson, S. Scarpino, B. Kerr, J. Rodrigues, J. Diaz-Colunga, and K. Kabengele for helpful interactions on the manuscript topic. The authors acknowledge support from the National Institutes of Health grant R35GM136354 (M.D.S.), R35GM147107 (R.F.G.), R01AI168166 (M.D.S. and C.B.O.), and the National Science Foundation’s Division of Environmental Biology Award Number 2142719 (C.B.O.). The authors would also like to thank the Martin Luther King Jr Visiting Professors and Scholars Program at the Massachusetts Institute of Technology for support (C.B.O.). Lastly, the authors would like to thank the organizers and participants in the 2022 workshop entitled “Reimagining the Central Dogma” at The Foundations Institute, University of California, Santa Barbara, where ideas relevant to this manuscript were discussed.

## Author contributions

Project conception: C.B.O. Collected and analyzed data: C.B.O., R.F.G. Interpreted and integrated data: C.B.O., R.F.G., E.I.S, M.D.S. Supervision: C.B.O., E.I.S., M.D.S. Writing: C.B.O., R.F.G., E.I.S., M.D.S.

## Data availability

Data and code area available: https://github.com/OgPlexus/subspace1

## Footnotes

1 We might contrive a new term that describes how epistatic interactions between mutations manifest across different traits. “Pleio-stasis” is a natural chimeric term that captures the essence of pleiotropy and epistasis. We have not used such a term in the main text of this study, however, as it may confuse readers.

