## Supplemental Text for "Epistasis meets pleiotropy in shaping biophysical protein subspaces associated with antimicrobial resistance"

### Supplemental Information

In the supplemental information, we provide the following:

- Data and analysis and code
- **Figure S1**: Rank orders of the TEM-1/TEM-50 alleles corresponding to those in Figure 2.
- **Table S1**: Sequence identity matrix for the three species analyzed in this study (*E. coli*, *L. grayi*, *C. muridarum*)

**Data and code**. Data and code can be found on `https://github.com/OgPlexus/subspace1`

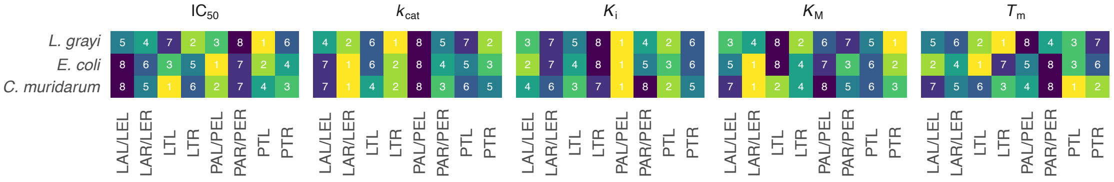


Figure S1: Rank orders of the TEM-1/TEM-50 alleles corresponding to those in Figure 2. We have utilized binary notation for this representation. As outlined in the METHODS section, binary notation corresponds to different alleles across the different species. For *E. coli* and *L. grayi*: 000 (PAL), 100 (LAL), 001(PAR), 010 (PTL), 100 (LAL) 011 (PTR),101 (LAR), 110(LTL), 111(LTR). For *C. muridarum*: 000 (PEL), 100 (LEL), 001 (PER), 010 (PTL), 100(LEL), 011 (PTR), 101 (LER), 110(LTL), 111(LTR).

| | *E. coli* | *L. grayi* | *C. muridarum* |
|---|---|---|---|
| *E. coli* | | | |
| *L. grayi* | 36 | | |
| *C. muridarum* | 27 | 23 | |

Table S1: Sequence identity matrix. Blocked out sections represent spaces with redundant information.
